# Can good broiler flock welfare prevent colonization by *Campylobacter*?

**DOI:** 10.1101/2021.03.30.437710

**Authors:** Thomas Rawson, Frances M. Colles, Adrian L. Smith, Marian Stamp Dawkins, Michael B. Bonsall

## Abstract

Using data on rearing and welfare metrics of multiple commercial broiler flocks from the last ten years, we investigate how welfare measures such as hock burn, mortality, weight, and pododermatitis, among others, impact the likelihood of a flock becoming colonized by *Campylobacter*. Using both logistic regression and Bayesian networks, we show that, while some welfare metrics were weakly related to *Campylobacter* colonization, evidence could not be found to suggest that these metrics actively exacerbated *Campylobacter* colonization, rather that they were both symptoms of the same underlying cause. Instead, observed dependency on the management of the flock suggested that yet-undiscovered differences in rearing practise were the principal cause of both poor bird welfare and increased risk of *Campylobacter*, suggesting that action can be taken to improve both these factors simultaneously.

## INTRODUCTION

For several years campylobacteriosis has been the most frequently observed zoonotic disease in humans throughout the EU (Westrell et al., 2009), with poultry meat identified as a leading infection route (EFSA Panel on Biological Hazards (BIOHAZ), 2011). This acute form of food poisoning, characterised by diarrhea, fever, and abdominal pain, is estimated to affect 450,000 individuals a year in the UK, approximately ten percent of which result in hospitalisation (Strachan and Forbes, 2010). An investigation by Public Health England into the extent of *Campylobacter* within the poultry industry revealed that seventy-three percent of supermarket chicken carcasses were found to contain *Campylobacter* and seven percent of the outer packaging was similarly contaminated (Jorgensen F, Madden RH, Arnold E, Charlett A, Elviss NC, 2015). This considerable public health burden posed by *Campylobacter* spp. represents an estimated £50 million annual economic cost to the UK (Tam and O’Brien, 2016).

Given the extent to which *Campylobacter* dominates commercial chicken flocks, attempting to reduce the proliferation of the pathogen at farm level would have significant impacts in reducing disease incidence in humans. Once *Campylobacter* is first identified within a broiler flock (chickens grown specifically for their meat), colonization of all birds occurs very rapidly (Evans and Sayers, 2000). In experimental studies, it can take only a single week for an entire flock to become infected following the introduction of a single infected bird (Stern et al., 2001). This speed of proliferation makes identifying the initial point of entry of *Campylobacter* into a flock challenging, and has resulted in a focus on preventative measures.

To-date, the poultry industry has largely focused upon on-farm biosecurity measures (Fraser et al., 2010; Gibbens et al., 2001), such as boot-dips and improved cleaning of housing. However, little impact in reducing incidence has been achieved with these measures (Hermans et al., 2011). As such, research has instead turned to a broad array of preventative measures (Ghareeb et al., 2013), such as treatment of food and water (Peh et al., 2020), probiotics (Saint-Cyr et al., 2016), and bacteriophage therapy (El-Shibiny et al., 2009). Such measures have thus far had mixed, and at times contradictory, success.

One area of research still greatly overlooked is the role of bird welfare in the emergence of *Campylobacter* within a flock, both as a potential indicator of *Campylobacter* colonization, and as a driving factor. *Campylobacter* spp. were long considered to be commensal within broiler chickens, but recent studies have begun to suggest they may be pathogenic under some circumstances (Humphrey et al., 2014; Wigley, 2015). Some welfare measures in the past have been observed to correlate with changes in the gut microbiota and immune response of birds, such as stocking density (Gomes et al., 2014; Guardia et al., 2011), food withdrawal, and heat stress (Burkholder et al., 2008). More directly, lesions on the footpad and arthritis have been shown to be strong predictors of *Campylobacter* prevalence (Alpigiani et al., 2017), further supporting findings that flock movement patterns and behaviour can also accurately predict *Campylobacter* prevalence (Colles et al., 2016). Our own previous mathematical modelling studies have highlighted the potential for stocking density (Rawson et al., 2019) to impact the population dynamics of *Campylobacter* within a flock, and have also shown that the colonization status of an entire flock is greatly impacted by the most susceptible birds within the flock (Rawson et al., 2020), suggesting that attention to individual birds must not be overlooked.

This study investigates the relationship between multiple welfare indicators on *Campylobacter* prevalence in flocks using two different mathematical modelling approaches. We firstly employ a logistic regression analysis to test for direct relationships between *Campylobacter* colonization and predictor variables, such as weight, mortality, and hock burn incidence. While this methodology has long served as a useful tool for highlighting potential relationships between variables, it cannot elucidate the exact mechanism of such a relationship, nor how these relationships interact with one another. We combine our logistic regression with a Bayesian network analysis to demonstrate the network of conditional dependencies between variables, to investigate more precisely how variables affect and impact each other. In combination with the logistic regression analysis, we are able to posit where welfare directly increases the likelihood of *Campylobacter* colonization, or to what extent infection by this bacteria is a symptom of the same root cause.

The greatest challenges to welfare-focused studies is ensuring a broad collection of data from varied sources, and using easily reproducible metrics. Studies utilising welfare concepts such as the ‘Welfare Quality^®^’ (De Jong et al., 2016) or the ‘five freedoms’ (Iannetti et al., 2020) are useful, but can be difficult to recreate due to differences in individual assessment. To this end, this study uses data spanning six years from multiple farms, logging reproducible metrics, such as temperature, flock parent age, pododermatitis rates, and flock size, amongst others.

## MATERIALS AND METHODS

### Data

Data was provided across six years (2010 to 2015) from multiple farms throughout the UK. Each data point represents a flock of broilers, listing multiple welfare parameters and rearing information, as well as a measure of whether the flock tested positive for *Campylobacter*. All variables measured for flocks are detailed and defined below:

- ***Company*** - A two-factor categorical variable, depicting whether the flock is overseen by company “1” or “2”. This variable will also therefore capture differences in company-specific rearing methodologies not represented by our current list of predictor variables.
- ***Farm*** - A categorical variable, further delineating the *Company* measure, detailing which farm the flock was located at, so as to investigate trends unique to certain locations.
- ***Number placed*** - A numerical variable describing how many broilers made up the flock. While modelling studies have primarily implicated stocking density as a high *Campylobacter* risk factor, the total flock population may also increase the likelihood of initial flock inoculation (Rawson et al., 2019).
- ***Date placed*** - The date the flock was first placed into the house. *Campylobacter* is well reported to show seasonal trends, with the warmer, summer months seeing flocks test positive for *Campylobacter* more frequently (Djennad et al., 2019; Nylen et al., 2002).
- ***Breed*** - A three-factor variable describing the breed of broilers grown. Two commercial breeds of broiler were investigated, with flocks comprised of either: Breed A, Breed B, or a mixture of Breed A & B. Both breeds refer to the breeding companies, each with many different genetic lines of broiler. Host-bird genetics have been shown to impact *Campylobacter* prevalence (Babacan et al., 2020; Psifidi et al., 2021; Stern et al., 1990), hence the consideration of the genetic line of the flock.
- ***Number of parent flocks*** - The number of parent flocks the broiler flock was sourced from. While the possibility of vertical transmission of *Campylobacter* is still debated, the hypothesis is that a greater number of parent flocks could increase the number of *Campylobacter* sequence types (and thus phenotypic specialisations) that a flock is exposed to at hatch (Petersen et al., 2001).
- ***Mean parent age*** - The average age (in weeks) of all parent flocks sourced from. Parent age has been shown to impact egg weight and embryo weight (Shanawany, 1984), and thus could potentially impact the general health of the chick.
- ***7/14/21/28/35/Total mortality percentage*** - Six different variables, describing the percentage of the flock that had died after *x* days.
- ***Pododermatitis percentage*** - What percentage of the flock suffered from pododermatitis; inflammation and ulcers on the footpad and toes. This was measured post-mortem by abattoir staff.
- ***Hock burn percentage*** - What percentage of the flock suffered from hock burn; areas where ammonia from the waste of other birds has burned through the skin of the leg. This was measured post-mortem by abattoir staff.
- ***7/14/21/28/35/Final day weight*** - Six variables showing the mean weight of the flock, in grams, at weekly intervals.
- ***Maximum/minimum temperature*** - A variable describing the maximum and minimum recorded *external* temperature, in degrees centigrade, for the time the flock was housed, as sourced from historical records for from the Met Office for the nearest weather station.
- ***Campylobacter 21/28/35 days*** - A two-factor variable depicting whether a flock was found to be positive or negative for *Campylobacter* after 21/28/35 days. This was sampled via fabric boot swabs in the flock house at 21/28/35 days. In addition, fresh faecal samples were collected concurrently on day 28. *Campylobacter* prevalence was then tested for in all samples via culture methods. Full details of this methodology are given in Colles et al. (2016).

A total of 212 flocks were monitored, however not all variables could be measured for all flocks due to the practical difficulties in obtaining all measures from farms. As such there is some degree of missing data across all variables, most notably that only 149 of these flocks have a final record of *Campylobacter* infection status. Before incorporating this data into a mathematical model, we must consider the detail of data available given the absence of some variables for some flocks. To ensure the maximum number of flocks are able to be used in model fitting, a balance must be found between filtering out variables to increase data availability, while not overly limiting the number of variables investigated. We detail these decisions below.

### Data *Cleaning*

Before beginning the regression analysis, we clean and simplify our data to aid interpretation. The *Campylobacter* variables across time points 21, 28, and 35 days were simplified to a single variable that reads as positive if a flock was recorded positive on any of the three dates recorded, and negative if the flock was reported negative on all of the measured dates provided. This was to increase the data availability, as some flocks were only measured on certain dates. There were six instances of a flock being recorded as negative after previously testing positive. These six instances were cases where the faecal samples taken on day 28 tested positive, but the boot swab on day 35 tested negative. It was considered appropriate to rely on the more targeted faecal sample for these six cases.

The *Date placed* variable was converted to a 4-level factor variable, denoting the season that the flock was reared in. This was done as date is known to have a non-linear effect on *Campylobacter* prevalence (Jorgensen et al., 2011), with incidence in both flocks and humans more frequently observed in the UK summer compared to the winter (Louis et al., 2005). It is this effect that we wish to investigate as opposed to variation between years. Season classification is partitioned by the dates December 1st, March 1st, June 1st, and September 1st, aligning with the meteorological seasons, which more accurately capture temperature variation than the astronomical seasons classification.

Regression analysis requires that the explanatory variables be independent of the response variable (and each other) otherwise predictive power is weakened across all dependent descriptor variables. In some cases, parameters of the linear model then become indeterminate due to the high degree of multicollinearity. For example, the *7/14/21/28/35/Total mortality percentage* variables are, as expected, all highly correlated with one another, hence we use only the *28-day mortality percentage* measure, as this is the one that most data was available for. We do the same for the average bird weight variables. Likewise, the *Company* variable was removed for the logistic regression, as it is heavily correlated with the *Farm* variable (companies do not share farms), however the *Farm* variable was also then found to have very strong correlation with the *Number placed* variable. For this reason the *Farm* variable is also removed, as *Number placed* is a preferred metric of interest. Similarly, we use only the *Minimum temperature*, and not the *Maximum temperature*, or the *Date placed*, as these three are strongly correlated. By reducing the number of model predictors, the generalised variance-inflation factors (GVIF) (Fox and Monette, 1992) of all variables are less than 3, far less than the commonly-used threshold of 10.

Finally, the data was filtered to remove any flocks with missing values for the explanatory variables under consideration. 84 data points remained for the final mathematical model. Flocks with missing data were later utilised for the parameter learning stage. A summary table of all variables considered in the final model is presented in Tables 1 and 2.

**Table 1.** Factor variable summaries

| Variable | Factor level | Total | *Campylobacter* positive | *Campylobacter* negative |
| --- | --- | --- | --- | --- |
| *Company* | 1 | 49 | 22 | 27 |
|  | 2 | 35 | 20 | 15 |
| *Farm* * | C1-F1 | 49 | 22 | 27 |
|  | C2-F1 | 15 | 7 | 8 |
|  | C2-F2 | 11 | 7 | 4 |
|  | C2-F3 | 9 | 6 | 3 |
| *Breed* | A | 32 | 20 | 12 |
|  | B | 48 | 21 | 27 |
|  | A & B | 4 | 1 | 3 |
| *No. Parents* | 1 | 39 | 23 | 16 |
|  | 2 | 25 | 13 | 12 |
|  | 3 | 15 | 5 | 10 |
|  | 4 | 5 | 1 | 4 |
| *Date placed* | Spring | 24 | 19 | 5 |
|  | Summer | 18 | 17 | 1 |
|  | Autumn | 16 | 1 | 15 |
|  | Winter | 26 | 5 | 21 |

\* Company 1 has only one farm ‘C1-F1’. Company 2 has three farms; ‘C2-F1’, ‘C2-F2’, ‘C2-F3’.

**Table 2.** Continuous variable summaries

| Variable | Mean | Standard Deviation |
| --- | --- | --- |
| *Number placed* | 27,639 | 7,283 |
| *Mean parent age* | 39.13 | 9.82 |
| *28-day mortality percentage* | 3.87 | 1.40 |
| *Pododermatitis percent* | 57.62 | 28.48 |
| *Hock burn percent* | 20.00 | 18.95 |
| *28-day average bird weight* | 1419.6 | 83.2 |
| *Minimum temperature* | 6.59 | 3.70 |

### Logistic Regression

Multiple logistic regression is an adaptation of multiple linear regression for instances where the response variable of interest is a two-factor binary output (*Y* ∈ {0,1}), in our case where a flock is either *Campylobacter* negative or positive. A multiple linear regression model structures the response variable, *Y*, as a linear predictor of a set of explanatory variables, *X_i_*, like so;

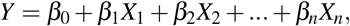

for *n* variables, and where *β_i_* are the coefficients to be determined. A logistic regression instead models *p* = *P*(*Y* = 1), theprobability that *Y* = 1, as:

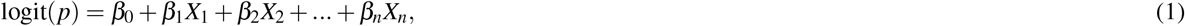

where logit() is the log-odds ratio 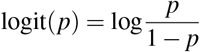, which ensures that *p* is bounded between 0 and 1. To model the impact of factor variables with *m* levels, we use treatment contrasts; *m* − 1 distinct descriptor variables within the model. For example, consider a simplified model which investigated the impact of breed alone on the probability of a flock being colonized by *Campylobacter* (*p*). *Breed* has three factor levels; ‘Breed A’, ‘Breed B’, and ‘Breed A & B’, and therefore the logistic regression model would be:

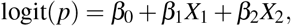

where Breed A, is represented by *X*_1_ = *X*_2_ = 0, Breed B by *X*_1_ = 1, *X*_2_ = 0, and the mixture of Breed A & B by *X*_2_ = 1, *X*_1_ = 0.

Nine explanatory variables were used for the final maximal logistic regression fit: *Number placed, Breed, Mean parent age, Number of parent flocks, 28-day mortality percentage, Pododermatitis percentage, Hock burn percentage, 28-day average weight*, and *Minimum temperature*. After initially fitting the maximal model of nine explanatory variables, a step wise simplification is then performed, removing the least significant term iteratively to finally reach the minimal adequate model: a model composed of only statistically significant explanatory variables. The model was fit using the glm package in R, which fits the model via iteratively reweighted least squares (IWLS). All code is made freely available at osf.io/pb62g/.

### Bayesian network

Bayesian networks are probabilistic graphical models that display the network of conditional dependencies between a collection of variables. Each variable in the model is visually represented as a node, with directed edges, called ‘arcs’, between nodes representing a directly dependent relationship. *A* → *B* indicates that *B* depends on *A*. Since arcs are directed, there is a cause-and-effect (from-and-to) relationship between variables. A node with an arc directed towards another node is called a ‘parent’ node to the respective ‘child’ node. Each node’s output is then explicitly detailed by a probability distribution that is dependent on any and all parent variables. This highlights the two greatest strengths of Bayesian networks as tools to investigate relationships between variables: firstly, the Markov property imposed by the network of conditional dependencies, means that the global probability distribution of the system can be expressed as a far smaller product of dependent probabilities. As such, a large and complicated probabilistic system can be simplified by knowledge of how some variables do or do not influence one another. Secondly, these types of models provide a straightforward way of visually conveying how certain explanatory variables influence (or do not influence) each other, something that would otherwise require the analysis of a large variety of logistic regression models, and could easily overlook certain dependencies. As a result of this architecture, “cycles” are by definition not allowed within a Bayesian network, meaning a path cannot be drawn from any node back to itself. Such a structure is called a directed acyclic graph (DAG). We provide a short example below to understand how such networks are calculated, but greater insight can be found in Nagarajan et al. (2013).

Calculating a Bayesian network model is separated into two tasks. Firstly, structural learning: learning the network model of dependencies (i.e. identifying all arcs in the system), followed by parameter learning: finding the specific parameters of probability distributions linking parent to child nodes. Consider an example of a dataset of three discrete variables in a broiler flock we wish to explore: *Mortality* (low, average, high), *Age* (young, adult, old), and *Feather condition* (good, average, poor). We start by learning the structure between these three variables. Many algorithms and approaches exist for finding the structure of a Bayesian network (Bouchaala et al., 2010), however within this paper we utilise the hill-climbing algorithm (Bouckaert, 1995), a score-based structure learning algorithm. The algorithm starts with a randomly chosen graph (though usually the empty graph made up of no arcs), and calculates a network score that ascertains how effectively such a graph describes the data. It then iteratively adds, removes and reverses one arc at a time, altering the global probability distribution via the introduction (or removal) of a dependency, selecting the alteration that increases the network score the most. This process is then repeated until no further improvement can be found. Multiple network scores can be used, but we use the Bayesian information criterion (BIC) (Bhat and Kumar, 2010), a variation on the traditional likelihood function. After using this algorithm on our example data, we discover the “best” network as being the network of two arcs shown in Figure 1.

**Figure 1.**
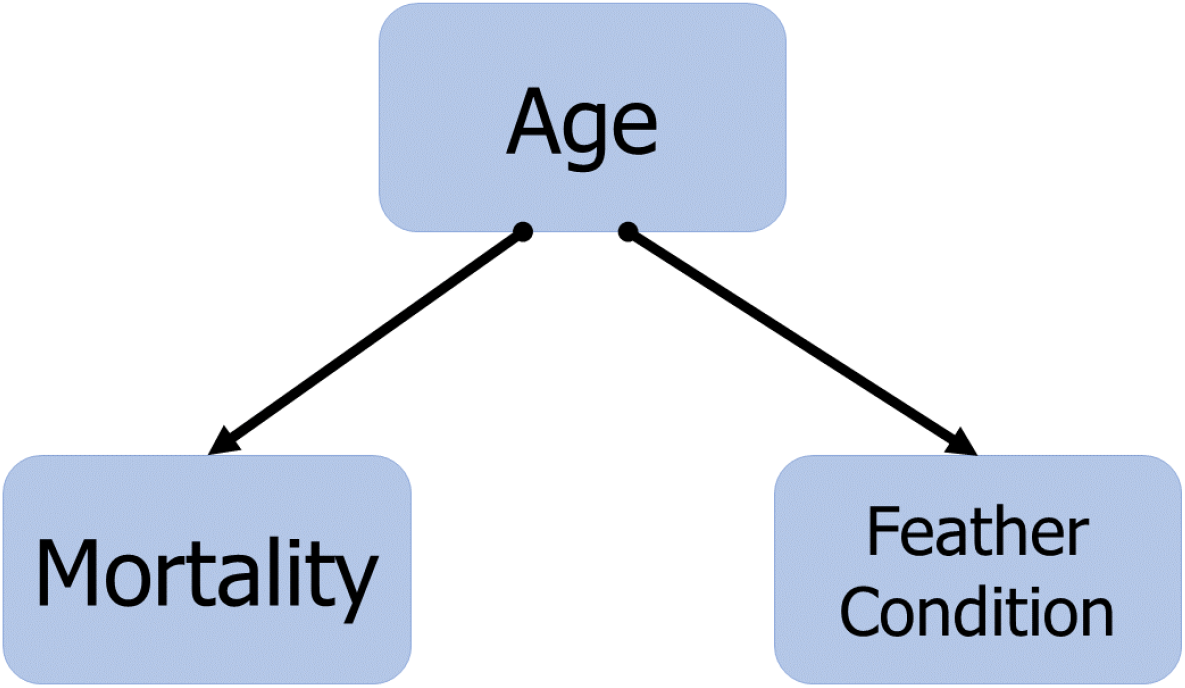
Bayesian network for the example problem posed. Two conditionally dependent relationships are found, from *Age* to *Mortality* and from *Age* to *Feather condition*. This example relationship was demonstrated by Comin et al. (2019).

We see from Figure 1 that *Age* is a parent variable to both *Mortality* and *Feather condition*. This indicates that, from this imagined example data, *Age* directly informs the mortality rate of a bird and the feather condition of the bird (this result was directly demonstrated by Comin et al. (2019)). An important insight gained from this network analysis would be that *Mortality* and *Feather condition* would likely be found to be correlated via a logistic regression analysis. However, *Feather condition* itself does not affect *Mortality*, rather they are both impacted by the same direct cause; *Age*. This illustrative example shows the objective of the Bayesian network approach to our particular question; what **causes** *Campylobacter* to colonize a flock, rather than just what is correlated with *Campylobacter* colonization. Another advantage of such a model, means that inference can be made even with missing data. The network of Figure 1 presents a structure whereby the mortality of a bird can be predicted with data on their feather condition, as this gives important indication of what the age of the bird may be. In Bayesian terms, this informs our prior belief as to the age of the bird, thus impacting our posterior belief as to the mortality of the bird. In contrast, the logistic regression approach would require an assumption on the age of such an individual, usually the mean of the training data, but no such requirement exists for Bayesian networks. Note, however, that if the age of a broiler is known, the prediction of their relative mortality rate is not improved by further information on their feather condition, as mortality is found to be predicted by age alone.

Note also from Figure 1 the mathematical advantage of such a network for expressing the joint probability distribution of the system. By definition the arcs indicate that *P*(*Age, Mortality, Feather Condition*) can be expressed as

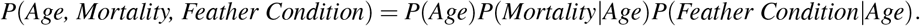

Since each variable has three factor levels, this reduces a distribution of 27 (3^3^) parameters, to 21, where each arc indicates that the child variable is modeled by a multinomial distribution dependent on the parent variable.

Indeed, for the second step, parameter estimation, we treat each node as being described by a multinomial distribution, and fit these using two separate well-known techniques, the maximum likelihood estimator (MLE), and a nested Bayesian approach, using uninformative uniform priors. See Appendix 1 for a brief introduction to Bayesian statistical inference.

A further benefit to a Bayesian network model is that we do not need to test for multicollinearity, which required us to remove several variables from consideration in the logistic regression, as structure learning specifically investigates these inter-variable correlations. As such we are able to include *Company*, *Farm*, and *Date placed* within our Bayesian network model. We also include the *7-day bird weight*, and *7-day mortality percentage* variables, alongside the 28-day measures, to serve both as a sanity check (we would expect these two variables to be linked), but also to increase the predictive power of the model, so inference could be made on the *Campylobacter* status of a flock from the 7-day as well as 28-day measures. This decision did however reduce the number of available training data from 84 to 81.

All of these introduced methodologies are implemented using the bnlearn package in R (Scutari, 2009), and all code used in the model analysis is provided at osf.io/pb62g.

### Discretisation

While we have displayed the many inherent strengths of Bayesian network models, one considerable weakness is the implementation of models consisting of both discrete and continuous variables. While methodologies exist for the assessment of such “hybrid” Bayesian networks (Scutari and Denis, 2014), the approaches are considerably more computationally demanding, and require a greater amount of data to give a robust fit to a Bayesian network. Given the comparatively smaller size of our training data (*n* = 84), we instead take the commonly used route of discretisation, whereby our continuous variables are converted into discrete bins. Of the many approaches to discretisation, a wide-ranging comparison by Kohavi and Sahami (1996) found the best approach to be the supervised, entropy-based Minimal Description Length (MDL) (Fayyad and Irani, 1993) method, whereby each variable is discretised based upon its informative potential on a variable of interest. This approach was undertaken on our data, in relation to the *Campylobacter* variable, using the `FSelectorRcpp` package in R. However, only *Minimum temperature* was found to be able to discretised in such a way (foreshadowing our later results). As such, we instead used a quantile binning (equal-frequency) approach, to separate out our continuous variables into three bins of equal frequency, and confirming against the histograms for each variable that no obvious separation was missed. These bin intervals are provided in Table 3.

**Table 3.** Discretisation intervals of continuous variables for the Bayesian network model

| Variable | Bin 1 Intervals | Bin 2 Intervals | Bin 3 Intervals |
| --- | --- | --- | --- |
| *Number placed* | [11770, 22000] | (22000, 33503.3] | (33503.3, 34650] |
| *Mean parent age* | [25, 32] | (32, 44.78] | (44.78, 58] |
| *7-day mortality percentage* | [0.65, 1.21] | (1.21, 1.81] | (1.81, 7.26] |
| *28-day mortality percentage* | [1.97, 3.06667] | (3.06667, 4.05667] | (4.05667, 9.61] |
| *Pododermatitis percent* | [1, 42.6667] | (42.6667, 76] | (76, 95] |
| *Hock burn percent* | [0, 10] | (10, 20] | (20, 90] |
| *7-day average bird weight* | [144, 170.667] | (170.667, 181] | (181, 213] |
| *28-day average bird weight* | [1138, 1388.33] | (1388.33, 1475.33] | (1475.33, 1565] |
| *Minimum temperature* | [1.3, 4] | (4, 8.7] | (8.7, 13.8] |

### Banned Arcs

To both aid the structure learning process, and to disallow erroneous network structures, we also introduce a list of banned arcs, defining all arcs which are not to be considered by the algorithm, based on logical reasoning. For example, we do not allow any arcs directed towards the *Company* variable, as this is clearly not affected by any other variables. While the company that a flock belongs to may in turn affect the mean parent bird age for example, it is illogical to say that the mean parent bird age could affect which company the flock is managed by. *Company* is a variable that is predetermined before the flock even hatches, and as such cannot be influenced by factors that occur during the lifespan of the flock. A full list of these banned illogical arcs is provided with all associated code in the online repository.

## RESULTS

### Logistic Regression

The results of the logistic regression for the minimal adequate model are presented in Table 4, alongside a variety of model evaluation metrics. Appendix 2 shows the analysis of the original maximal model comprised of all explanatory variables, and describes the reduction steps taken to reach the minimal adequate model.

**Table 4.** Logistic regression analysis of the minimal adequate model for 84 broiler flocks using the `glm` function in R.

| Predictor | $\beta$ (Estimate) | SE $\beta$ | Wald-test z-score | *p* | $e^{\beta}$ (odds ratio) |
| --- | --- | --- | --- | --- | --- |
| (Intercept) | -1.490 | 0.621 | -2.398 | **0.0165** | NA |
| *Breed* (1 = Breed B, 0 = Other) | -1.474 | 0.628 | -2.348 | **0.0189** | 0.225 |
| *Hock burn percentage* | -0.041 | 0.0189 | -2.165 | **0.0304** | 0.960 |
| *Minimum temperature* | 0.469 | 0.105 | 4.474 | **$\ll 0.0001$** | 1.599 |
| Test | | | $\chi^2$ | *p* | |
| Overall model evaluation |  |  |  |  |  |
| Likelihood ratio test <sup>1</sup> | | | 35.972 | $7.59 \times 10^{-8}$ | |
| Likelihood ratio test <sup>2</sup> |  |  | 7.776 | 0.557 |  |
| Goodness of fit |  |  |  |  |  |
| McFadden's $R^2$ | 0.309 | | | | |
| Cox & Snell's $R^2$ | 0.348 | | | | |

<sup>1</sup> Compared against null model.<sup>2</sup> Compared against maximal model.

Three variables were found to be statistically significant in relation to the *Campylobacter* status of a flock via the Wald-test: *Breed, Hock burn percentage*, and *Minimum temperature*. Note that for the minimum adequate model, while Breed B flocks were found to be statistically significantly different to Breed A birds with relation to *Campylobacter* incidence, the mixed breed flocks were not found to differ from Breed A flocks. As such, the ‘Breed A’ and ‘Breed A & B’ flocks were collapsed into one variable for the minimal adequate model. Table 4 shows that flocks of Breed B birds were found to be 0.225 times as likely to test positive for *Campylobacter* than any other flock. *Hock burn percentage* was, unintuitively, found to have a negative correlation with *Campylobacter* colonization. *Minimum temperature* was very strongly correlated, with an odds ratio showing that an increase of 1 degree to the minimum recorded temperature corresponded with a flock being 1.599 times more likely to test positive for *Campylobacter*. The generalised variance-inflation factors (GVIF) (Fox and Monette, 1992) of all variables in the minimal adequate model was less than 2, and all variables of the maximal model (Appendix 2) had a GVIF of less than 3, far less than the commonly-used threshold of 10.

### Bayesian network

Figure 2 displays the final global network structure, fit using the hill-climbing algorithm, and with networks scored via BIC. This was run using the `bnlearn` package in R. The strength of individual arcs (as measured by BIC) is represented by arrow-width in Figure 2. Table 5 also explicitly provides these arc strength scores.

**Figure 2.**
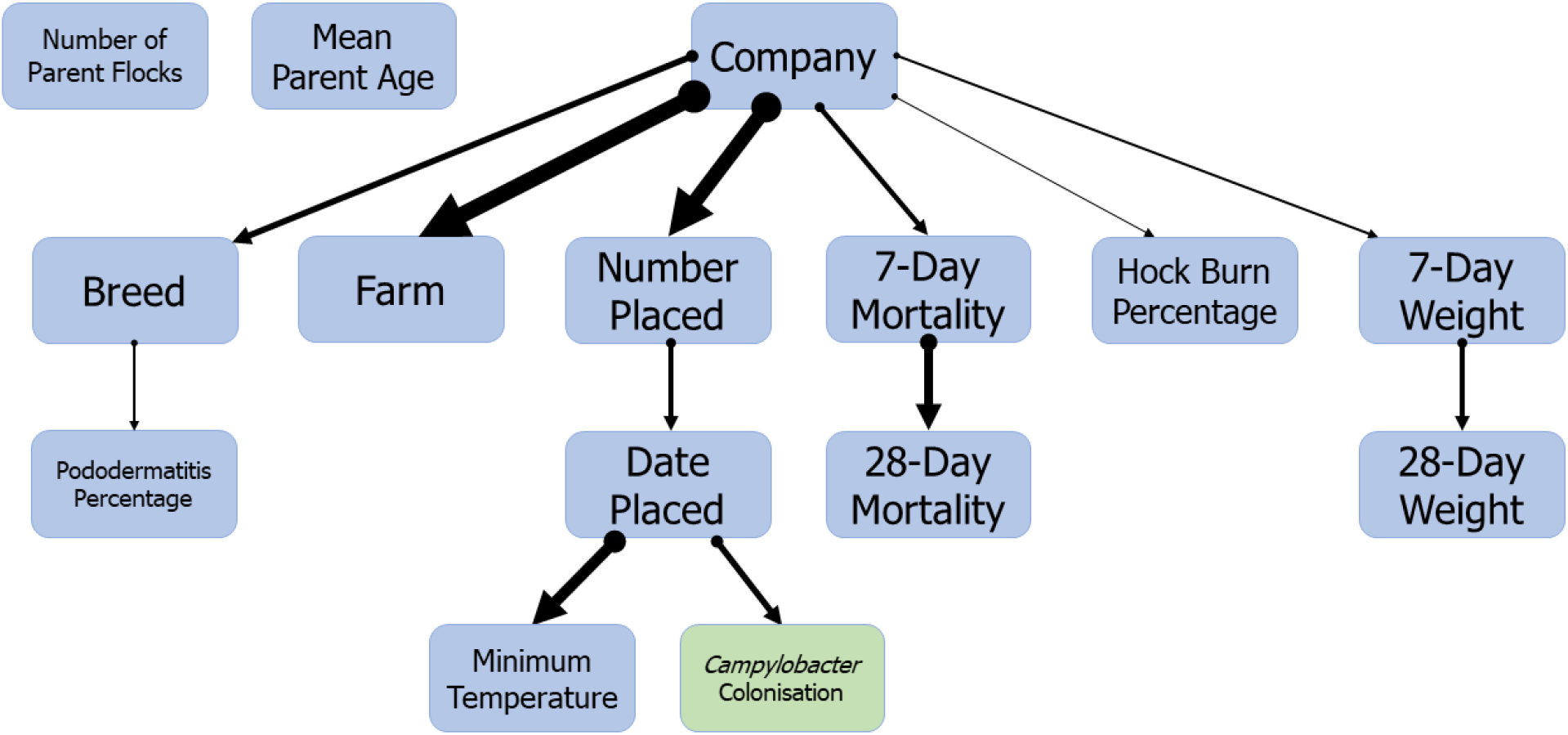
Bayesian network structure showing the interrelationships between multiple welfare and rearing practice factors in a flock of broilers. *Campylobacter* colonization is directly impacted by the season the flock is grown in. Structure was learned using a hill-climbing algorithm, and sampled networks scored using the Bayesian information criterion (BIC). Arrow-width indicates arc strength as scored by BIC, the values of which are given in Table 5.

**Table 5.** Arc strengths of the Bayesian network shown in Figure 2. Arc strength is measured by Bayesian information criterion (BIC), where a lower value indicators a stronger link.

| Parent | Child | Arc Strength |
| --- | --- | --- |
| *Company* | *Breed* | -16.656 |
| *Company* | *Farm* | -47.756 |
| *Company* | *Number placed* | -43.069 |
| *Company* | *7-day mortality percentage* | -11.975 |
| *Company* | *Hock burn percentage* | -0.234 |
| *Company* | *7-day weight* | -3.644 |
| *Breed* | *Pododermatitis percentage* | -2.236 |
| *Number placed* | *Date placed* | -6.272 |
| *7-day mortality percentage* | *28-day mortality percentage* | -20.876 |
| *7-day weight* | *28-day weight* | -6.031 |
| *Date placed* | *Minimum temperature* | -29.040 |
| *Date placed* | *Campylobacter colonization* | -17.246 |

To test the significance of the fit structure, structure learning was also performed with a tabu search algorithm, and by introducing random network restarts into the hill-climbing algorithm (10, 100, and 1000 random restarts were all performed), all of which resulted in the same network structure. We also performed a hill-climbing structure learning algorithm using the logarithm of the Bayesian Dirichlet equivalent score (BDE) (Castelo and Siebes, 2000), as opposed to the BIC, a Bayesian-based score equivalent to the Dirichlet posterior density (and initialised with uniform priors). This scoring metric resulted in a very similar network structure which we present in Appendix 3. The only differences were that, (i) *Hock burn percentage* no longer had *Company* as a parent node, meaning it was unconnected to any other node. (ii) *Minimum temperature* had an additional arc from itself to *Campylobacter* colonization, suggesting that *Campylobacter* could also be impacted by temperature variation throughout the season; and finally, (iii) an additional arc was introduced from *Breed* to *Number placed*, simply representing that flocks of Breed B birds were larger in size than flocks of Breed A birds.

Other results to be noted from Figure 2 is that neither the number of different parent flocks that a broiler flock was born from, nor the mean age of these parent was found to have any correlation to any other variable. *Pododermatitis* was interestingly found to be influenced by the *Breed* of broiler comprising the flock. We also see that many variables are directly influenced by the *Company* variable, suggesting that many observed differences are due to, yet unobserved, differences between management practise.

Figure 2 shows that the season (*Date placed*) in which a flock is reared is the sole parent node to *Campylobacter* status. This means that *Date placed* alone captures the uncertainty and probabilistic distribution of whether or not a flock is likely to test positive for *Campylobacter*. This means that while data on the number of birds in the flock (*Number placed*) can inform whether or not a flock is *Campylobacter* positive, this data is superfluous when one has knowledge of the *Date placed*. The conditional probability table for *Campylobacter* colonization is given in Table 6. These model parameters can be fit either via maximum likelihood estimators (MLE) or through Bayesian inference. Model parameters via both methods are provided in Table 6. Note that one advantage of the Bayesian inference method is that this approach can learn parameters from data containing missing values. Hence while the MLE parameters are fit from the 84 data points used in structure learning, the Bayesian inference method uses 114 data points, incorporating those that were removed from structure learning due to missing values.

**Table 6.** Conditional probability table for *Campylobacter* colonization status. We present values calculated by Bayesian inference using uniform priors, and an equivalent sample size of 10. Below that we present values calculated via a maximum likelihood estimator.

| Bayesian Inference |  |  |  |  |
| --- | --- | --- | --- | --- |
| *Date placed* |  |  |  |  |
| *Campylobacter* colonization | Spring | Summer | Autumn | Winter |
| Negative | 0.273 | 0.071 | 0.878 | 0.802 |
| Positive | 0.727 | 0.929 | 0.122 | 0.198 |

| Maximum Likelihood Estimator |  |  |  |  |
| --- | --- | --- | --- | --- |
| *Date placed* |  |  |  |  |
| *Campylobacter* colonization | Spring | Summer | Autumn | Winter |
| Negative | 0.217 | 0.063 | 0.937 | 0.808 |
| Positive | 0.783 | 0.937 | 0.063 | 0.192 |

Conditional probability tables for *Campylobacter* dependent on all other variables, assuming the absence of data on any other variable, are provided in Appendix 4.

## DISCUSSION

Here, through a combination of both logistic regression and Bayesian network analysis, we have investigated the interrelationships between a selection of welfare and rearing practice explanatory variables for multiple commercial broiler flocks. At the inception of this work, our hypothesis was that poor welfare indicators such as low weight and hock burn, among others, would result in an increased risk of colonization by *Campylobacter* due to poor health compromising the immune response of birds in the flock (Humphrey, 2006). Social stress (Mohamed and Hanson, 1980), heat stress (Burkholder et al., 2008), and overcrowding stress (Gomes et al., 2014), have all been shown to increase susceptibility to disease in chickens by compromising the immune response (Heckert et al., 2002; Hirakawa et al., 2020), and in many cases have been correlated with increased risk of colonization with *Salmonella* (Alhenaky et al., 2017; Gomes et al., 2014). As such it was initially assumed that similar measures may increase incidence of *Campylobacter* in broiler flocks. While our work has revealed some level of correlation between poor welfare metrics and *Campylobacter* incidence (see the conditional probabilities of Appendix 4), these relationships were not found to be statistically significant via a logistic regression model, and our Bayesian network model suggests that poor bird welfare, as judged by the measures used here, is not in fact a cause of increased *Campylobacter* colonization. Despite this, our model reveals many yet-unconsidered relationships between rearing variables, provides evidence against multiple existing hypotheses, and highlights multiple promising new lines of enquiry towards identifying the source of *Campylobacter* colonization in commercial poultry flocks.

Our logistic regression analysis, shown in Table 4, identified three statistically significant explanatory variables; *Breed*, *Minimum temperature* and *Hock burn percentage*, with *p* values of 0.0189, 7.69 × 10^-6^ and 0.0304 respectively. Seasonal variation in *Campylobacter* incidence has long been observed in broiler flocks (Jorgensen et al., 2011; Louis et al., 2005), with minimum/maximum temperature and sunshine hours significantly correlated with both the incidence and total bacterial load found in chicken flocks (Wallace et al., 1997). The warmer summer months see greater *Campylobacter* prevalence, yet despite the large body of research confirming this phenomenon, the precise mechanism for this increase remains unclear. While the growth rate of *Campylobacter* is found to vary in relation to temperature (Doyle and Roman, 1981), the minimum temperature required for *Campylobacter* survival is estimated to be around 30 degrees centigrade, somewhat precluding the impact of UK seasonal temperatures. Previous studies have suggested that the seasonal increase of flies (Hald et al., 2004, 2007), rodents (Meerburg and Kijlstra, 2007), and wild birds (Colles et al., 2008) as vectors of *Campylobacter* transmission may be responsible, while seasonal patterns in country-wide clonal complex incidence potentially point to genetic adaptation to seasonal trends (Jorgensen et al., 2011). Investigating this trend in human incidence of campylobacteriosis, Djennad et al. (2019) conducted a rigorous statistical assessment of spatial and weather factors, concluding that the correlation between incidence and temperature was “ likely to be indirect”. Our above results reach the same conclusion for broiler colonization. While our logistic regression shows the strong correlation between temperature and *Campylobacter* colonization, our Bayesian network analysis shows in Figure 2 that the two variables are conditionally independent upon the date placed, i.e. the correlation is indirect.

Footpad dermatitis, commonly referred to as ‘hock burn’, the dark discolouration and ulceration of the lower leg of birds, was also found to be statistically significantly correlated with *Campylobacter* prevalence, however this relationship was curiously found to be negatively correlated. These painful lesions are considered a sign of poor bird welfare, usually caused by litter unsuitably saturated with chicken waste. As such, the suggestion that more instances of hock burn in a flock are linked with less cases of *Campylobacter* is surprising, considering that the bacteria are transmitted via the faecal-oral route. One hypothesis is that the presence of *Campylobacter* may in turn limit colonisation of the flock by more pathogenic bacteria that could more easily trigger diarrhoea within a host-bird, thus impacting the litter quality and the resulting development of hock burn. Alternatively this relationship may be an artifact of how the *Hock burn percentage* variable was defined. Namely it was recorded as the cross-sectional prevalence of any signs of hockburn in the flock (Dawkins et al., 2017). In short, it is a measure of how many birds showed signs of hock burn, and not a measure of the extremity of these burns. Bull et al. (2008) observed this same effect, whereby the flock-wide presence of hock burn was generally higher in *Campylobacter* negative flocks, however the number of birds in the flock rejected from consumption due to extreme cases of hock burn was positively correlated with rates of *Campylobacter* colonization. Figure 2 also concludes that this correlation between *Campylobacter* colonization and hock burn prevalence is conditionally independent upon the managing company.

The Bayesian network structure displayed in Figure 2 reveals a wide variety of insight into the various interrelationships of the included variables. Firstly we see that the number of parent flocks a broiler flock is sourced from, and the mean age of these parent flocks, had no meaningful impact on association with any other variable. The feasibility of vertical transmission of *Campylobacter* from parent to broiler flock is still frequently discussed in the literature, and the inclusion of this variable was based upon the hypothesis that a greater number of parent flocks may challenge a broiler flock with a greater genotypic variety of *Campylobacter* isolates (Petersen et al., 2001). Parent age has also been shown to influence egg weight and embryo weight of chicks (Shanawany, 1984). Given the potential importance of maternal antibodies in suppressing *Campylobacter* in the first few weeks of age (Rawson et al., 2019), parent age could potentially impact the likelihood of *Campylobacter* colonization. Our results however indicate that factors relating to the parent flock have no effect on any of the metrics considered in this study.

The logistic regression analysis of the minimum adequate model found statistical significance in the relationship between *Breed* and *Campylobacter* colonization, where flocks of Breed A birds were more frequently observed to become colonized than Breed B birds. Caffrey et al. (2021) recently identified a correlation between breed and *Campylobacter*, with flocks comprised of Cobb birds, or a mixture of Cobb and Ross birds 4.75 times more likely to test positive for fluoroquinolone resistant *Campylobacter jejuni* than flocks comprised of just Ross birds. Further to this, Cobb birds have been found to be more frequently colonized by *Campylobacter* than Hubbard birds by Babacan et al. (2020), however they were unable to separate this association from other rearing factors such as age-of-slaughter. Our Bayesian network analysis, similar to the hock burn conclusions, was unable to detect any direct arc of causation between *Breed* and *Campylobacter* colonization, suggesting that the breed of chicken is indicative of the company managing the flock, and not necessarily an indicator of a breed-specific susceptibility. Host-bird genetics have however previously been shown to cause differences in host-resistance to *Campylobacter* challenge (Connell et al., 2013; Li et al., 2008; Stern et al., 1990), with such resistances shown to be inheritable under experimental conditions Boyd et al. (2005). Further linking breed and welfare measures, Humphrey et al. (2014) found that faster-growing breeds of broiler showed evidence of prolonged inflammation in the intestines in response to *Campylobacter jejuni*, suggesting that the impact of breed is yet a plausible route of further study. An interesting relationship observed in Figure 2 was the implication of *Breed* as a determinant of the prevalence of pododermatitis, with flocks of Breed A birds more frequently displaying heavy incidence of pododermatitis. No study to our knowledge has directly investigated this supposed relationship in broilers, however one study in turkeys found no correlation between breed and pododermatitis (Clark et al., 2002). Pododermatitis has previously been shown to be associated with a poor-nutrient diet (Nagaraj et al., 2007), hence the hypothesis that this factor could correlate with general gut health and/or the composition of the gut microbiome.

The primary conclusion of our work, as shown in Figure 2, was that our network of variables was closely related by yet-unobserved factors concealed within the *Company* variable. *Company* was found to be a parent variable to six factors; *Breed, Farm, Number placed, 7-day mortality, Hock burn percentage*, and *7-day weight*. This indicates that these six factors significantly vary, due to which of the two companies considered within this study they are managed by. This suggests that choices made within the complex decision network relating to the rearing of these flocks, encompassing factors such as diet, water provision, housing, thinning protocols, cleaning regimens, antibiotic usage, and stocking density among others (Sibanda et al., 2018), will have the significant potential to both decrease incidence of *Campylobacter* and may simultaneously improve the welfare of the flock. While disappointing to not ascertain the primary root cause of increased *Campylobacter* prevalence from within our considered set of variables, the work has revealed a key network of dependencies within commonly recorded and studied metrics. While far from the first study to examine the contributions of multiple health factors towards *Campylobacter* colonization (Babacan et al., 2020; Frosth et al., 2020; Humphery et al., 1993; Rushton et al., 2009), our work is the first, to our knowledge, to utilise the powerful methodologies underlying Bayesian network analysis in studying the spread of *Campylobacter*. Such approaches, in combination with more traditional logistic regression analyses, greatly increase the descriptive power of gathered datasets, and it is our hope that this work will help expedite their adoption throughout the field of *Campylobacter* risk management. Bayesian networks have had some early success already in specifically implicating welfare measures with specific housing variables (Comin et al., 2019), we now further our attempts to identify the variables that exacerbate the spread of *Campylobacter*.

This study illustrates the need to investigate, more thoroughly, management decisions in the broiler industry, so as to reduce *Campylobacter* incidence whilst improving bird health and welfare, to provide the consumer with a better product whilst reducing the impact of campylobacteriosis on human health.

## Author contributions statement

F.M.C. performed the microbiological sampling. All authors interpreted the results. T.R., M.B.B. and M.S.D. conceived the study. T.R. built the models and wrote all associated code. T.R. wrote the manuscript. M.S.D., F.M.C., and M.B.B. supervised the project. All authors reviewed the manuscript.

## A Appendices

### A.1 Appendix 1 - Bayesian Statistics

This brief section aims to convey the basic principles of Bayesian statistics, and familiarise the reader with the terminology that is be used throughout the manuscript. For an in-depth explanation, we recommend the text by Kruschke (2014).

Bayesian statistics is derived wholly from the relationship defined by Bayes’ theorem,

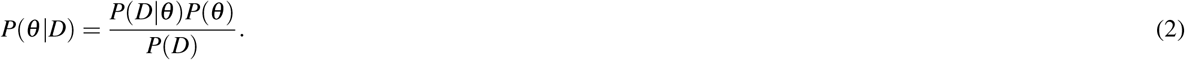

If we consider *θ* as some statistical parameter we wish to infer, and *D* as some data informing the parameter, then equation (1) expresses that the probability distribution for our value of *θ*, given our dataset (*P*(*θ*|*D*)), is proportional to the **likelihood** of such data (*P*(*D*| *θ*)) multiplied by the probability distribution of *θ* free of any data (*P*(*θ*)).

One starts with a **prior** probabilistic understanding of the values *θ*, often informed by expert opinion, and by utilising relevant data, D, we update our belief in the values *θ* may take, producing a new **posterior** distribution. Mnemonically, if we wished to calculate the probability that a flipped coin will land heads up, we may have a **prior** belief that the coin is fair. However, upon observing a data set of 5 coin flips, all of which produced heads, we may update our **posterior** belief to reflect that the coin may be biased.

The analytical difficulty in this calculation lies in computing *P*(*D*) = ∫*P*(*D*|*θ*)*P*(*θ*)d*θ*, which is often near impossible for realistically complex models. Fortunately modern computing power enables us to efficiently estimate our posterior distributions through algorithms such as Gibbs sampling and other Metropolis-Hastings schemes.

Hierarchical systems represent multi-variable models where some parameters depend on other parameters. Returning to the example of a coin flip, say the probability of heads (*θ*) is dependent on the factory in which the coin was minted. The probability that a coin was from a certain factory (*ω*) will then inform our value of (*θ*). Expressed mathematically, equation (1) now becomes:

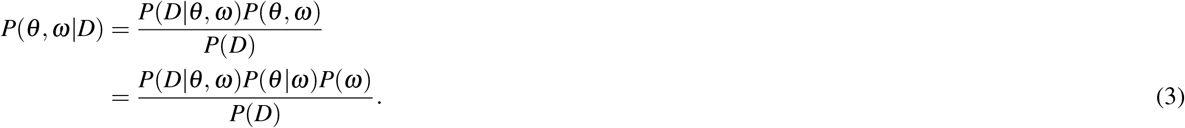

This means that a prior distribution is only required for *ω*, as this distribution will directly inform our **conditional prior** of *θ*, via our model formulation. As such, when provided with data on coin flips from multiple coins from different factories, we obtain a posterior probability distribution of which factory a coin has come from, and the resulting probability of a coin flip resulting in heads. This structure of conditional independence means that data relating specifically to one parameter can still help inform the posterior of all other dependent variables, a key advantage of Bayesian inference.

### A.2 Appendix 2 - Logistic Regression maximal model to minimal adequate model

First we present the logistic regression analysis of the maximal model:

**Table 7.** Logistic regression analysis of the maximal model.

| Predictor | $\beta$ (Estimate) | SE $\beta$ | Wald-test $z$-score | $p$ | $e^{\beta}$ (odds ratio) |
| --- | --- | --- | --- | --- | --- |
| (Intercept) | 8.958 | 7.00 | 1.28 | 0.2006 | NA |
| *Number placed* | $-5.435 \times 10^{-5}$ | $6.928 \times 10^{-5}$ | -0.785 | 0.433 | 0.999 |
| *Breed* (1 = B, 0 = A) | -1.510 | 0.788 | -1.916 | 0.0554 | 0.221 |
| *Breed* (1 = A & B, 0 = A) | -1.213 | 1.815 | -0.668 | 0.504 | 0.297 |
| *Mean parent age* | 0.0255 | 0.0406 | 0.629 | 0.529 | 1.026 |
| *No. parent flocks* (1 = 2 flocks, 0 = 1 flock) | -0.456 | 0.755 | -0.604 | 0.546 | 0.638 |
| *No. parent flocks* (1 = 3 flocks, 0 = 1 flock) | -0.930 | 0.846 | -1.1 | 0.271 | 0.394 |
| *No. parent flocks* (1 = 4 flocks, 0 = 1 flock) | -1.372 | 1.519 | -0.903 | 0.367 | 0.254 |
| *28-day mortality percentage* | -0.339 | 0.318 | -1.065 | 0.287 | 0.713 |
| *Pododermatitis percentage* | -0.016 | 0.015 | -1.142 | 0.253 | 0.983 |
| *Hock burn percentage* | -0.0531 | 0.0240 | -2.215 | **0.0268** | 0.948 |
| *28-day average weight* | -0.0050 | 0.0047 | -1.070 | 0.284 | 0.995 |
| *Minimum temperature* | 0.512 | 0.135 | 3.809 | **0.0001** | 1.669 |
| Test | | | $\chi^2$ | $p$ | |
| Overall model evaluation * |  |  |  |  |  |
| Likelihood ratio test | | | 43.748 | $1.685 \times 10^{-5}$ | |
| Goodness of fit test |  |  |  |  |  |
| Hosmer-Lemeshow |  |  | 7.9779 | 0.436 |  |
| McFadden's $R^2$ | 0.376 | | | | |
| Cox & Snell's $R^2$ | 0.406 | | | | |

\* Compared against null model.

We then iteratively remove the least significant variables, until only significant variables remain. Firstly we remove *Mean parent age*.

**Table 8.** Logistic regression analysis of the maximal model with *Mean parent age* removed.

| Predictor | $\beta$ (Estimate) | SE $\beta$ | Wald-test z-score | *p* |
| --- | --- | --- | --- | --- |
| (Intercept) | 8.887 | 6.863 | 1.295 | 0.195 |
| *Number placed* | $-7.137 \times 10^{-5}$ | $6.412 \times 10^{-5}$ | -1.113 | 0.195 |
| *Breed* (1 = B, 0 = A) | -1.382 | 0.755 | -1.832 | 0.067 |
| *Breed* (1 = A & B, 0 = A) | -1.155 | 1.806 | -0.639 | 0.523 |
| *No. parent flocks* (1 = 2 flocks, 0 = 1 flock) | -0.461 | 0.749 | -0.616 | 0.538 |
| *No. parent flocks* (1 = 3 flocks, 0 = 1 flock) | -0.973 | 0.850 | -1.144 | 0.253 |
| *No. parent flocks* (1 = 4 flocks, 0 = 1 flock) | -1.426 | 1.556 | -0.916 | 0.360 |
| *28-day mortality percentage* | -0.392 | 0.312 | -1.257 | 0.209 |
| *Pododermatitis percentage* | -0.017 | 0.015 | -1.160 | 0.246 |
| *Hock burn percentage* | -0.0527 | 0.0239 | -2.201 | **0.0277** |
| *28-day average weight* | -0.0038 | 0.0041 | -0.917 | 0.359 |
| *Minimum temperature* | 0.497 | 0.128 | 3.883 | **0.0001** |
| Test | | | $\chi^2$ | *p* |
| Overall model evaluation |  |  |  |  |
| Likelihood ratio test <sup>1</sup> | | | 47.16 | $2.014 \times 10^{-6}$ |
| Likelihood ratio test <sup>2</sup> |  |  | 0.4033 | 0.5254 |
| Goodness of fit test |  |  |  |  |
| Hosmer-Lemeshow |  |  | 11.131 | 0.1944 |
| McFadden's $R^2$ | 0.372 | | | |
| Cox & Snell's $R^2$ | 0.403 | | | |

<sup>1</sup> Compared against null model.
<sup>2</sup> Compared against maximal model.

Next we remove Number of *parent flocks*.

**Table 9.**
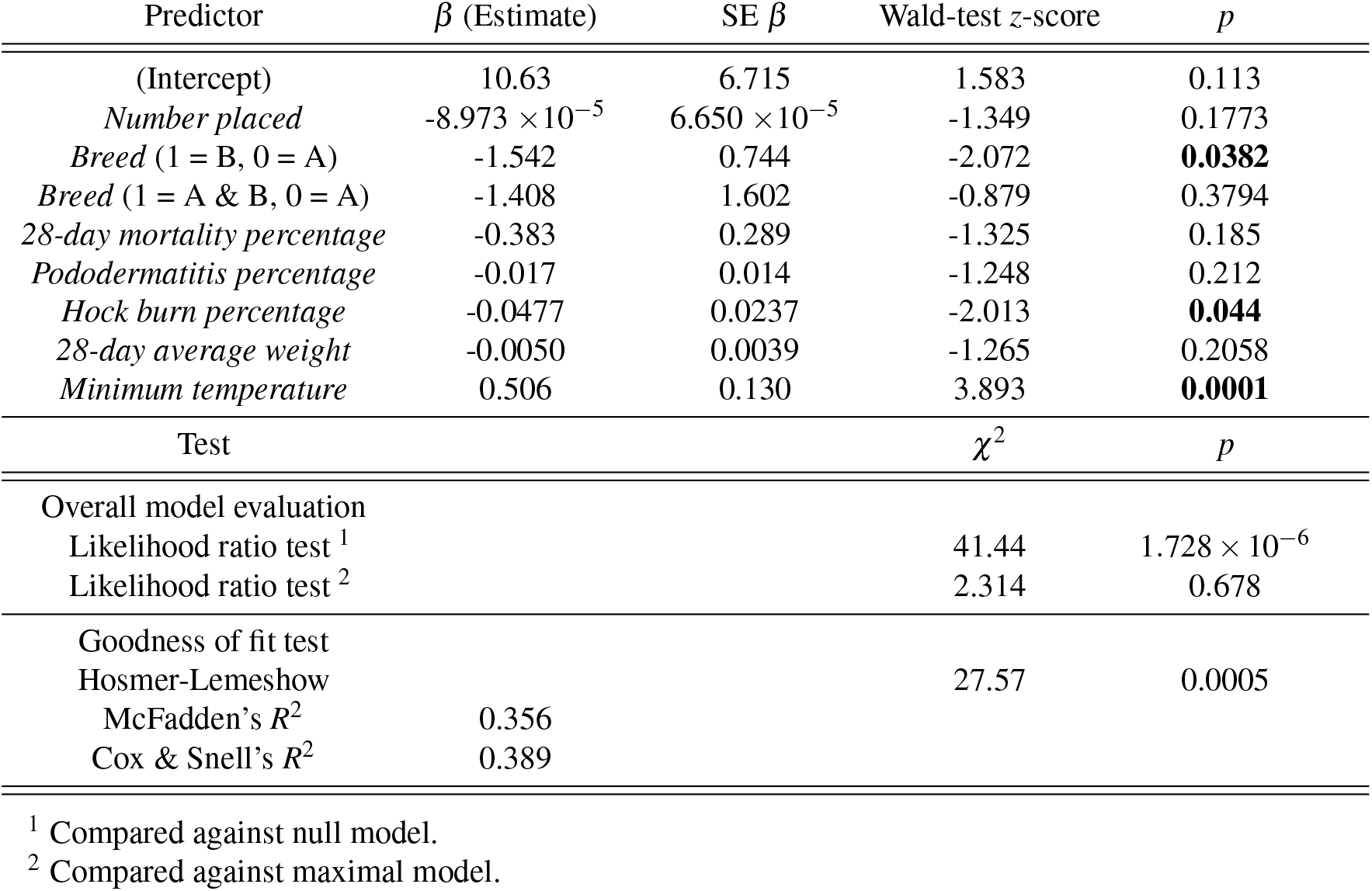
Logistic regression analysis of the maximal model with *Number of parent flocks* removed.

| Predictor | $\beta$ (Estimate) | SE $\beta$ | Wald-test z-score | *p* |
| --- | --- | --- | --- | --- |
| (Intercept) | 10.63 | 6.715 | 1.583 | 0.113 |
| *Number placed* | $-8.973 \times 10^{-5}$ | $6.650 \times 10^{-5}$ | -1.349 | 0.1773 |
| *Breed* (1 = B, 0 = A) | -1.542 | 0.744 | -2.072 | **0.0382** |
| *Breed* (1 = A & B, 0 = A) | -1.408 | 1.602 | -0.879 | 0.3794 |
| *28-day mortality percentage* | -0.383 | 0.289 | -1.325 | 0.185 |
| *Pododermatitis percentage* | -0.017 | 0.014 | -1.248 | 0.212 |
| *Hock burn percentage* | -0.0477 | 0.0237 | -2.013 | **0.044** |
| *28-day average weight* | -0.0050 | 0.0039 | -1.265 | 0.2058 |
| *Minimum temperature* | 0.506 | 0.130 | 3.893 | **0.0001** |
| Test | | | $\chi^2$ | *p* |
| Overall model evaluation |  |  |  |  |
| Likelihood ratio test <sup>1</sup> | | | 41.44 | $1.728 \times 10^{-6}$ |
| Likelihood ratio test <sup>2</sup> |  |  | 2.314 | 0.678 |
| Goodness of fit test |  |  |  |  |
| Hosmer-Lemeshow |  |  | 27.57 | 0.0005 |
| McFadden's $R^2$ | 0.356 | | | |
| Cox & Snell's $R^2$ | 0.389 | | | |

<sup>1</sup> Compared against null model.
<sup>2</sup> Compared against maximal model.

Next we remove *Pododermatitis percentage*.

**Table 10.** Logistic regression analysis of the maximal model with *Pododermatitis percentage* removed.

| Predictor | $\beta$ (Estimate) | SE $\beta$ | Wald-test z-score | *p* |
| --- | --- | --- | --- | --- |
| (Intercept) | 8.086 | 6.309 | 1.282 | 0.200 |
| *Number placed* | $-6.77 \times 10^{-5}$ | $6.029 \times 10^{-5}$ | -1.123 | 0.2615 |
| *Breed* (1 = B, 0 = A) | -1.501 | 0.735 | -2.043 | **0.0410** |
| *Breed* (1 = A & B, 0 = A) | -0.647 | 1.456 | -0.444 | 0.657 |
| *28-day mortality percentage* | -0.366 | 0.283 | -1.295 | 0.195 |
| *Hock burn percentage* | -0.0519 | 0.0228 | -2.283 | **0.0224** |
| *28-day average weight* | -0.0046 | 0.0038 | -1.186 | 0.2356 |
| *Minimum temperature* | 0.540 | 0.129 | 4.192 | **0.00003** |
| Test | | | $\chi^2$ | *p* |
| Overall model evaluation |  |  |  |  |
| Likelihood ratio test <sup>1</sup> | | | 39.76 | $1.398 \times 10^{-6}$ |
| Likelihood ratio test <sup>2</sup> |  |  | 3.986 | 0.551 |
| Goodness of fit test |  |  |  |  |
| Hosmer-Lemeshow |  |  | 25.464 | 0.00130 |
| McFadden's $R^2$ | 0.341 | | | |
| Cox & Snell's $R^2$ | 0.377 | | | |

<sup>1</sup> Compared against null model.
<sup>2</sup> Compared against maximal model.

Next we remove *Number placed*.

**Table 11.** Logistic regression analysis of the maximal model with *Number placed* removed.

| Predictor | $\beta$ (Estimate) | SE $\beta$ | Wald-test z-score | *p* |
| --- | --- | --- | --- | --- |
| (Intercept) | 4.983 | 5.434 | 0.917 | 0.359 |
| *Breed* (1 = B, 0 = A) | -1.825 | 0.697 | -2.617 | **0.0089** |
| *Breed* (1 = A & B, 0 = A) | -1.267 | 1.358 | -0.933 | 0.351 |
| *28-day mortality percentage* | -0.228 | 0.238 | -0.956 | 0.339 |
| *Hock burn percentage* | -0.0513 | 0.0211 | -2.435 | **0.0149** |
| *28-day average weight* | -0.0037 | 0.0037 | -1.016 | 0.310 |
| *Minimum temperature* | 0.491 | 0.113 | 4.341 | **0.00001** |
| Test | | | $\chi^2$ | *p* |
| Overall model evaluation |  |  |  |  |
| Likelihood ratio test <sup>1</sup> | | | 38.512 | $8.921 \times 10^{-7}$ |
| Likelihood ratio test <sup>2</sup> |  |  | 5.237 | 0.514 |
| Goodness of fit test |  |  |  |  |
| Hosmer-Lemeshow |  |  | 22.82 | 0.0036 |
| McFadden's $R^2$ | 0.331 | | | |
| Cox & Snell's $R^2$ | 0.368 | | | |

<sup>1</sup> Compared against null model.<sup>2</sup> Compared against maximal model.

Next we remove *28-day mortality percentage*.

**Table 12.** Logistic regression analysis of the maximal model with *28-day mortality percentage* removed..

| Predictor | $\beta$ (Estimate) | SE $\beta$ | Wald-test z-score | $p$ |
| --- | --- | --- | --- | --- |
| (Intercept) | 3.835 | 5.266 | 0.728 | 0.466 |
| *Breed* (1 = B, 0 = A) | -1.639 | 0.652 | -2.513 | **0.0120** |
| *Breed* (1 = A & B, 0 = A) | -1.133 | 1.406 | -0.806 | 0.420 |
| *Hock burn percentage* | -0.045 | 0.019 | -2.33 | **0.0197** |
| *28-day average weight* | -0.0036 | 0.0037 | -0.978 | 0.328 |
| *Minimum temperature* | 0.471 | 0.108 | 4.344 | **0.00001** |
| Test | | | $\chi^2$ | $p$ |
| Overall model evaluation |  |  |  |  |
| Likelihood ratio test <sup>1</sup> | | | 37.59 | $4.553 \times 10^{-7}$ |
| Likelihood ratio test <sup>2</sup> |  |  | 6.155 | 0.522 |
| Goodness of fit test |  |  |  |  |
| Hosmer-Lemeshow |  |  | 23.16 | 0.0032 |
| McFadden's $R^2$ | 0.323 | | | |
| Cox & Snell's $R^2$ | 0.361 | | | |

<sup>1</sup> Compared against null model.
<sup>2</sup> Compared against maximal model.

Next we remove *28-day average weight*.

**Table 13.**
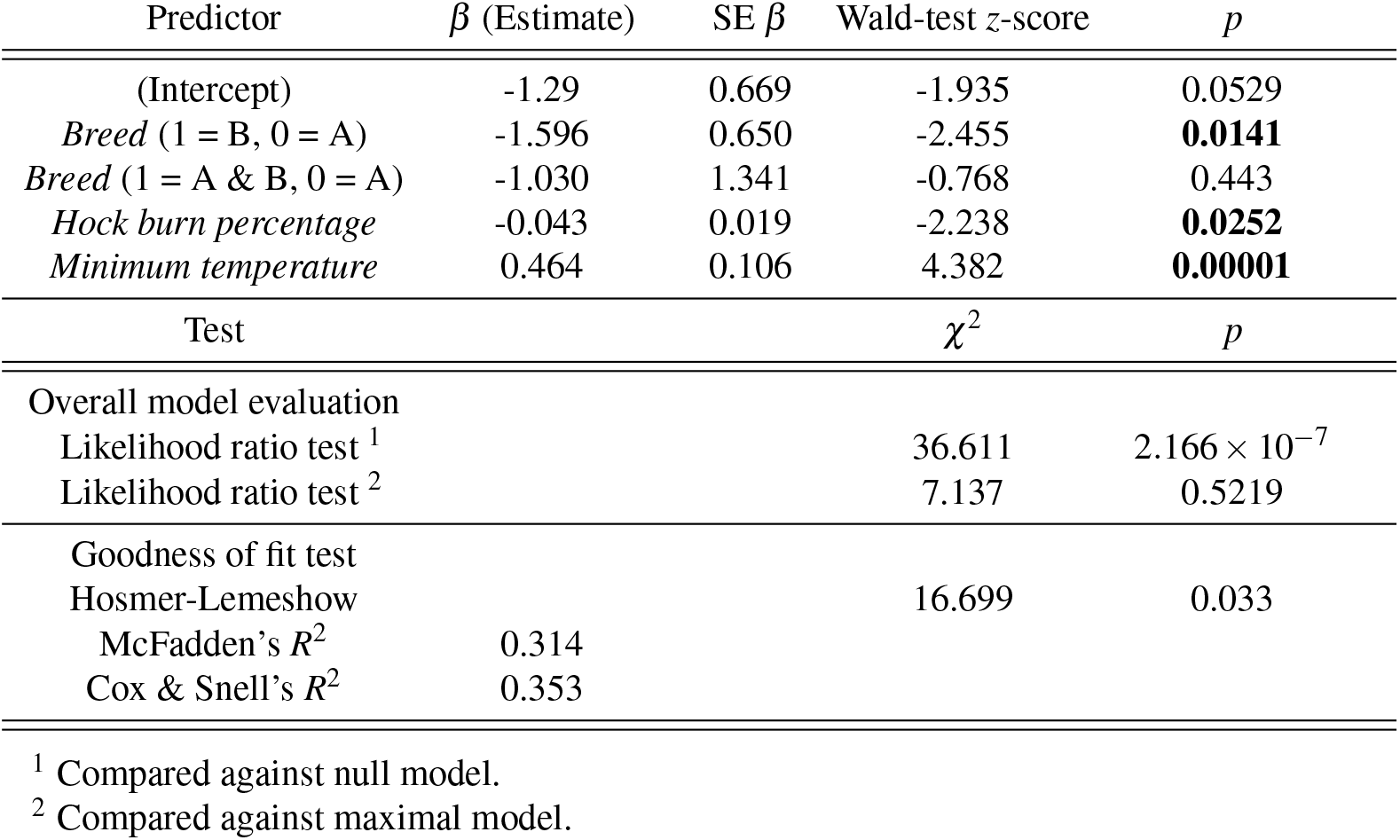
Logistic regression analysis of the maximal model with *28-day average weight* removed.

| Predictor | $\beta$ (Estimate) | SE $\beta$ | Wald-test z-score | *p* |
| --- | --- | --- | --- | --- |
| (Intercept) | -1.29 | 0.669 | -1.935 | 0.0529 |
| *Breed* (1 = B, 0 = A) | -1.596 | 0.650 | -2.455 | **0.0141** |
| *Breed* (1 = A & B, 0 = A) | -1.030 | 1.341 | -0.768 | 0.443 |
| *Hock burn percentage* | -0.043 | 0.019 | -2.238 | **0.0252** |
| *Minimum temperature* | 0.464 | 0.106 | 4.382 | **0.00001** |
| Test | | | $\chi^2$ | *p* |
| Overall model evaluation |  |  |  |  |
| Likelihood ratio test <sup>1</sup> | | | 36.611 | $2.166 \times 10^{-7}$ |
| Likelihood ratio test <sup>2</sup> |  |  | 7.137 | 0.5219 |
| Goodness of fit test |  |  |  |  |
| Hosmer-Lemeshow |  |  | 16.699 | 0.033 |
| McFadden's $R^2$ | 0.314 | | | |
| Cox & Snell's $R^2$ | 0.353 | | | |

<sup>1</sup> Compared against null model.
<sup>2</sup> Compared against maximal model.

The only remaining non-significant variable remains within the *Breed* variable. Table 13 shows that the factor ‘Breed A & B’ is not statistically significantly different in its predictive potential from Breed A birds. As such, we collapse the ‘Breed A’ and ‘Breed A & B’ factors together, to produce the final minimal adequate model as provided in Table 4 of the manuscript.

### A.3 Appendix 3 - Bayesian network structure with BDE scoring

Here we present the best-fit network structure when the hill-climbing algorithm is used with BDE scoring instead of BIC scoring.

**Figure 3.**
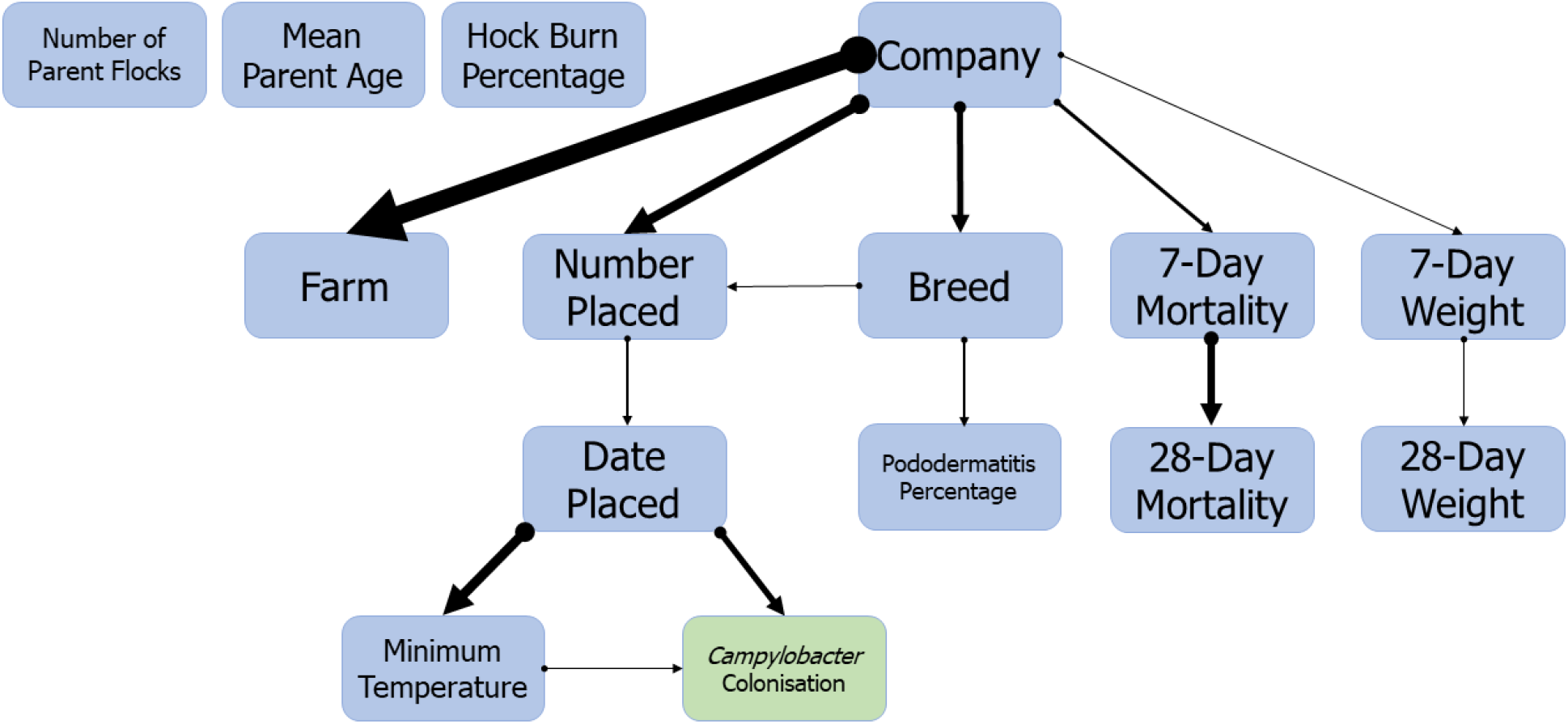
Bayesian network structure showing the interrelationships between multiple welfare and rearing practise factors in a flock of broilers. *Campylobacter* colonization is directly impacted by the season the flock is grown in. Structure was learned using a hill-climbing algorithm, and sampled networks scored using the Bayesian Dirichlet equivalent score (BDE). Arrow-width indicates arc strength as scored by BDE.

We provide the exact arc strengths as scored by BDE below.

**Table 14.**
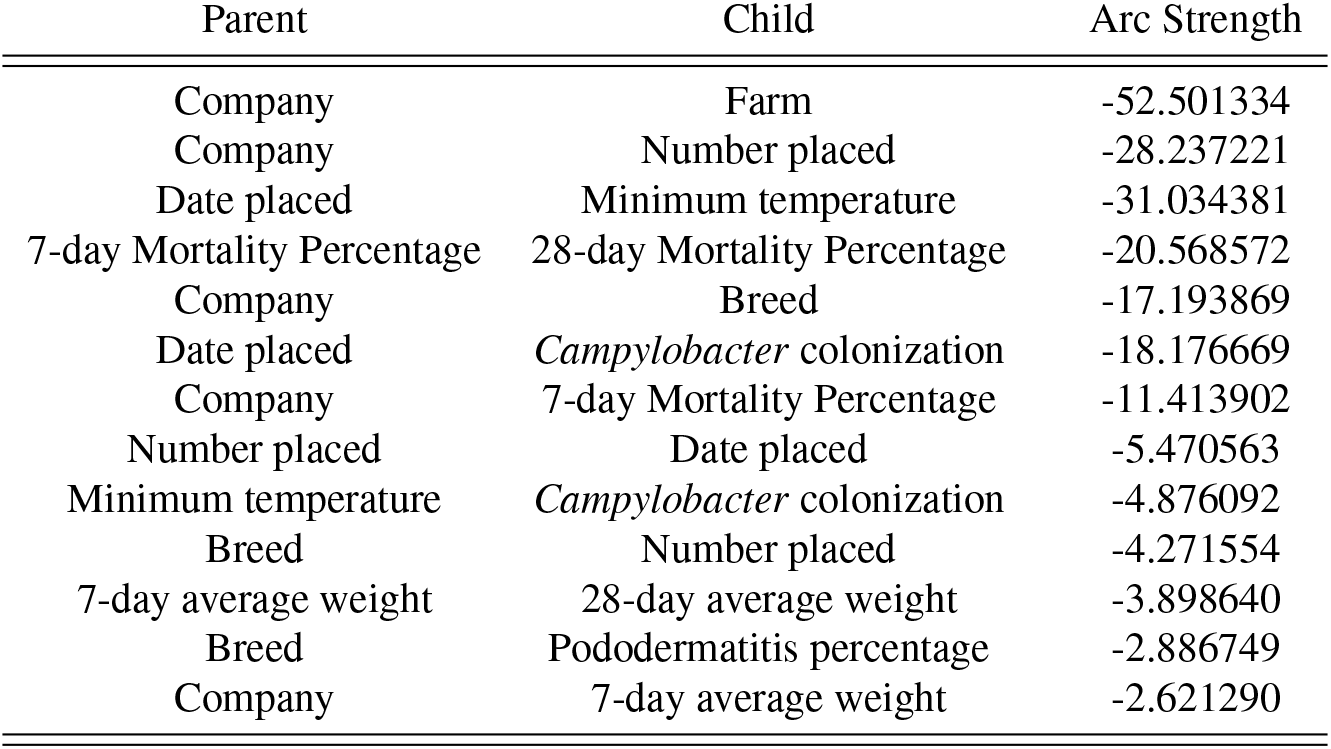
Arc strengths of the Bayesian network shown in Figure A.2.1. Arc strength is measured by Bayesian Dirichlet equivalent score (BDE), where a lower value indicators a stronger link.

| Parent | Child | Arc Strength |
| --- | --- | --- |
| Company | Farm | -52.501334 |
| Company | Number placed | -28.237221 |
| Date placed | Minimum temperature | -31.034381 |
| 7-day Mortality Percentage | 28-day Mortality Percentage | -20.568572 |
| Company | Breed | -17.193869 |
| Date placed | *Campylobacter* colonization | -18.176669 |
| Company | 7-day Mortality Percentage | -11.413902 |
| Number placed | Date placed | -5.470563 |
| Minimum temperature | *Campylobacter* colonization | -4.876092 |
| Breed | Number placed | -4.271554 |
| 7-day average weight | 28-day average weight | -3.898640 |
| Breed | Pododermatitis percentage | -2.886749 |
| Company | 7-day average weight | -2.621290 |

### A.4. Appendix 4 - *Campylobacter* conditional probability tables

The network structure shown in Figure 2 displays that the date placed alone captures the probabilistic distribution of whether or not a flock is colonized by *Campylobacter*. However, in the absence of data on the date placed, other predictor variables can inform our expectations of whether or not a flock will be *Campylobacter* positive. The following tables provide these conditional probabilities for *Campylobacter* colonization under the assumption that no data is known other than the variable displayed. The best fit parameters via both Bayesian inference and MLE are given. The Bayesian estimates are built from a larger dataset of 114 entries, 33 of which contain some missing data. We do not provide tables upon mean parent age or number of parent flocks, as these variables were found to be unassociated.

**Table 15.** Conditional probability table for *Campylobacter* colonization status, when data is only available on the Company variable. We present values calculated by Bayesian inference using uniform priors, and an equivalent sample size of 10. Below that we present values calculated via a maximum likelihood estimator.

| Bayesian Inference |  |  |
| --- | --- | --- |
|  | Company |  |
| *Campylobacter* colonization | 1 | 2 |
| Negative | 0.508 | 0.379 |
| Positive | 0.492 | 0.621 |
| Maximum Likelihood Estimator |  |  |
|  | Company |  |
| *Campylobacter* colonization | 1 | 2 |
| Negative | 0.576 | 0.430 |
| Positive | 0.424 | 0.570 |

**Table 16.** Conditional probability table for *Campylobacter* colonization status, when data is only available on the Farm variable. We present values calculated by Bayesian inference using uniform priors, and an equivalent sample size of 10. Below that we present values calculated via a maximum likelihood estimator.

| Bayesian Inference |  |  |  |  |
| --- | --- | --- | --- | --- |
|  | Farm |  |  |  |
| *Campylobacter* colonization | C1F1 | C2F1 | C2F2 | C2F3 |
| Negative | 0.505 | 0.386 | 0.387 | 0.388 |
| Positive | 0.495 | 0.614 | 0.613 | 0.612 |

| Maximum Likelihood Estimator |  |  |  |  |
| --- | --- | --- | --- | --- |
|  | Farm |  |  |  |
| *Campylobacter* colonization | C1F1 | C2F1 | C2F2 | C2F3 |
| Negative | 0.576 | 0.430 | 0.430 | 0.430 |
| Positive | 0.424 | 0.570 | 0.570 | 0.570 |

**Table 17.** Conditional probability table for *Campylobacter* colonization status, when data is only available on the ‘number placed’ variable. We present values calculated by Bayesian inference using uniform priors, and an equivalent sample size of 10. Below that we present values calculated via a maximum likelihood estimator.

| Bayesian Inference |  |  |  |
| --- | --- | --- | --- |
|  | Number placed |  |  |
| *Campylobacter* colonization | [11770,22000] | (22000,33503.3] | (33503.3,34650] |
| Negative | 0.362 | 0.533 | 0.533 |
| Positive | 0.638 | 0.467 | 0.467 |

| Maximum Likelihood Estimator |  |  |  |
| --- | --- | --- | --- |
|  | Number placed |  |  |
| *Campylobacter* colonization | [11770,22000] | (22000,33503.3] | (33503.3,34650] |
| Negative | 0.417 | 0.616 | 0.544 |
| Positive | 0.583 | 0.384 | 0.456 |

**Table 18.**
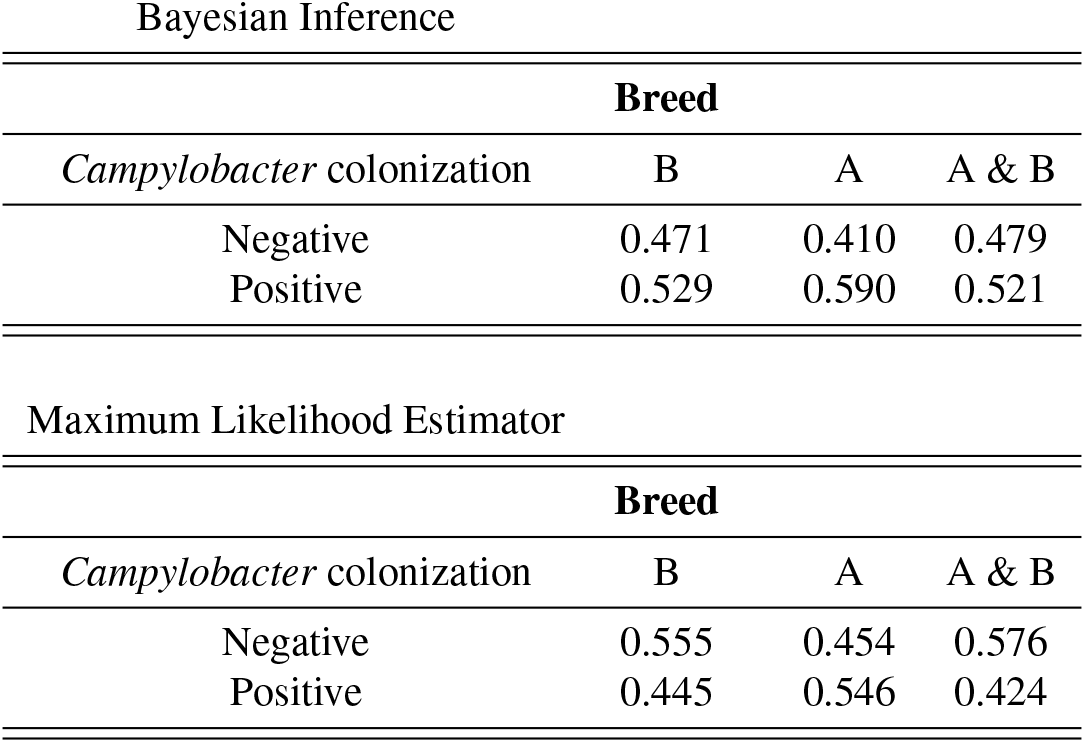
Conditional probability table for *Campylobacter* colonization status, when data is only available on the breed variable. We present values calculated by Bayesian inference using uniform priors, and an equivalent sample size of 10. Below that we present values calculated via a maximum likelihood estimator.

| Bayesian Inference |  |  |  |
| --- | --- | --- | --- |
| Breed |  |  |  |
| *Campylobacter* colonization | B | A | A & B |
| Negative | 0.471 | 0.410 | 0.479 |
| Positive | 0.529 | 0.590 | 0.521 |

| Maximum Likelihood Estimator |  |  |  |
| --- | --- | --- | --- |
| Breed |  |  |  |
| *Campylobacter* colonization | B | A | A & B |
| Negative | 0.555 | 0.454 | 0.576 |
| Positive | 0.445 | 0.546 | 0.424 |

**Table 19.** Conditional probability table for *Campylobacter* colonization status, when data is only available on the 7-day mortality variable. We present values calculated by Bayesian inference using uniform priors, and an equivalent sample size of 10. Below that we present values calculated via a maximum likelihood estimator.

| Bayesian Inference |  |  |  |
| --- | --- | --- | --- |
| 7-Day Mortality Percentage |  |  |  |
| *Campylobacter* colonization | [0.65,1.21] | (1.21,1.81] | (1.81,7.26] |
| Negative | 0.496 | 0.444 | 0.406 |
| Positive | 0.504 | 0.556 | 0.594 |

| Maximum Likelihood Estimator |  |  |  |
| --- | --- | --- | --- |
| 7-Day Mortality Percentage |  |  |  |
| *Campylobacter* colonization | [0.65,1.21] | (1.21,1.81] | (1.81,7.26] |
| Negative | 0.571 | 0.517 | 0.468 |
| Positive | 0.429 | 0.483 | 0.532 |

**Table 20.** Conditional probability table for *Campylobacter* colonization status, when data is only available on the 28-day mortality variable. We present values calculated by Bayesian inference using uniform priors, and an equivalent sample size of 10. Below that we present values calculated via a maximum likelihood estimator.

| Bayesian Inference |  |  |  |
| --- | --- | --- | --- |
| 28-Day Mortality Percentage |  |  |  |
| *Campylobacter* colonization | [1.97,3.06667] | (3.06667,4.05667] | (4.05667,9.61] |
| Negative | 0.480 | 0.445 | 0.419 |
| Positive | 0.520 | 0.555 | 0.581 |
| Maximum Likelihood Estimator |  |  |  |
| 28-Day Mortality Percentage |  |  |  |
| *Campylobacter* colonization | [1.97,3.06667] | (3.06667,4.05667] | (4.05667,9.61] |
| Negative | 0.557 | 0.518 | 0.481 |
| Positive | 0.443 | 0.482 | 0.519 |

**Table 21.** Conditional probability table for *Campylobacter* colonization status, when data is only available on the 7-day weight variable. We present values calculated by Bayesian inference using uniform priors, and an equivalent sample size of 10. Below that we present values calculated via a maximum likelihood estimator.

| Bayesian Inference |  |  |  |
| --- | --- | --- | --- |
| 7-Day Average Weight |  |  |  |
| *Campylobacter* colonization | [144,170.667] | (170.667,181] | (181,213] |
| Negative | 0.481 | 0.453 | 0.418 |
| Positive | 0.519 | 0.547 | 0.582 |
| Maximum Likelihood Estimator |  |  |  |
| 7-Day Average Weight |  |  |  |
| *Campylobacter* colonization | [144,170.667] | (170.667,181] | (181,213] |
| Negative | 0.555 | 0.522 | 0.479 |
| Positive | 0.445 | 0.478 | 0.521 |

**Table 22.** Conditional probability table for *Campylobacter* colonization status, when data is only available on the 28-day weight variable. We present values calculated by Bayesian inference using uniform priors, and an equivalent sample size of 10. Below that we present values calculated via a maximum likelihood estimator.

| Bayesian Inference |  |  |  |
| --- | --- | --- | --- |
| 28-Day Average Weight |  |  |  |
| *Campylobacter* colonization | [1138,1388.33] | (1388.33,1475.33] | (1475.33,1565] |
| Negative | 0.459 | 0.451 | 0.433 |
| Positive | 0.541 | 0.549 | 0.567 |
| Maximum Likelihood Estimator |  |  |  |
| 28-Day Average Weight |  |  |  |
| *Campylobacter* colonization | [1138,1388.33] | (1388.33,1475.33] | (1475.33,1565] |
| Negative | 0.538 | 0.522 | 0.496 |
| Positive | 0.462 | 0.478 | 0.504 |

**Table 23.** Conditional probability table for *Campylobacter* colonization status, when data is only available on the hock burn percentage variable. We present values calculated by Bayesian inference using uniform priors, and an equivalent sample size of 10. Below that we present values calculated via a maximum likelihood estimator.

| Bayesian Inference |  |  |  |
| --- | --- | --- | --- |
| Hock Burn Percentage |  |  |  |
| *Campylobacter* colonization | [0,10] | (10,20] | (20,90] |
| Negative | 0.438 | 0.447 | 0.452 |
| Positive | 0.562 | 0.553 | 0.548 |
| Maximum Likelihood Estimator |  |  |  |
| Hock Burn Percentage |  |  |  |
| *Campylobacter* colonization | [0,10] | (10,20] | (20,90] |
| Negative | 0.490 | 0.520 | 0.548 |
| Positive | 0.510 | 0.480 | 0.452 |

**Table 24.** Conditional probability table for *Campylobacter* colonization status, when data is only available on the pododermatitis percentage variable. We present values calculated by Bayesian inference using uniform priors, and an equivalent sample size of 10. Below that we present values calculated via a maximum likelihood estimator.

| Bayesian Inference |  |  |  |
| --- | --- | --- | --- |
| Pododermatitis Percentage |  |  |  |
| *Campylobacter* colonization | [1,42.6667] | (42.6667,76] | (76,95] |
| Negative | 0.448 | 0.457 | 0.436 |
| Positive | 0.552 | 0.543 | 0.564 |

| Maximum Likelihood Estimator |  |  |  |
| --- | --- | --- | --- |
| Pododermatitis Percentage |  |  |  |
| *Campylobacter* colonization | [1,42.6667] | (42.6667,76] | (76,95] |
| Negative | 0.524 | 0.537 | 0.490 |
| Positive | 0.476 | 0.463 | 0.510 |

**Table 25.** Conditional probability table for *Campylobacter* colonization status, when data is only available on the minimum temperature variable. We present values calculated by Bayesian inference using uniform priors, and an equivalent sample size of 10. Below that we present values calculated via a maximum likelihood estimator.

| Bayesian Inference |  |  |  |
| --- | --- | --- | --- |
| Minimum Temperature |  |  |  |
| *Campylobacter* colonization | [1.3,4] | (4,8.7] | (8.7,13.8] |
| Negative | 0.452 | 0.442 | 0.416 |
| Positive | 0.548 | 0.558 | 0.584 |

| Maximum Likelihood Estimator |  |  |  |
| --- | --- | --- | --- |
| Minimum Temperature |  |  |  |
| *Campylobacter* colonization | [1.3,4] | (4,8.7] | (8.7,13.8] |
| Negative | 0.694 | 0.465 | 0.313 |
| Positive | 0.306 | 0.535 | 0.687 |

